# The Role of Glycan Structures in Modulating GM-CSF Bioactivity: Insights from Glycoengineering

**DOI:** 10.64898/2026.02.17.706267

**Authors:** Ece Cagdas, Sarah Line Skovbakke, Judit Pina Agullet, Leo A. Dworkin, Giulia Scapin, Hooman Hefzi, Karoline Schousboe Fremming, Sanne Schoffhelen, Natalia Putkaradze, Bjørn Voldborg, Lise M. Grav, Lars K. Nielsen, Steffen Goletz, Nathan E. Lewis

## Abstract

Granulocyte-macrophage colony-stimulating factor (GM-CSF) is a glycoprotein cytokine with therapeutic potential in cancer and neutropenia treatment. While glycosylation of GM-CSF reduces immunogenicity and enhances serum bioavailability, it can also diminish receptor binding and bioactivity. Based on transcriptomic analysis of human T lymphocytes reported previously, GM-CSF–producing cells exhibit elevated expression of Alpha-1,6-Mannosylglycoprotein 6-Beta-N-Acetylglucosaminyltransferase (*MGAT5*), which encodes N-acetylglucosaminyltransferase V, an enzyme involved in N-glycan branching. Given this role of MGAT5 in glycosylation, we produced GM-CSF variants using glycoengineered Chinese hamster ovary cells to generate diverse glycoforms and assessed their bioactivity. Testing their activity on TF-1 cell proliferation, we found that decreases in GM-CSF *N*-glycan branching significantly suppressed its activity. These findings underscore the importance of glycosylation in modulating the efficacy and safety of GM-CSF-based therapeutics, suggesting that precise glycoengineering may be key to optimizing GM-CSF performance in clinical applications.

## Introduction

Cytokines often have important immunomodulatory properties, thus can be part of valuable therapies for autoimmune diseases and cancer. One such cytokine of therapeutic importance is GM-CSF. This glycosylated cytokine induces the maturation and proliferation of myeloid precursors ^1^. It can be produced from diverse types of cells, such as T cells, endothelial, epithelial, and fibroblast cells ^2^. It has been associated with a range of diseases and has been the subject of numerous clinical trials ^3,4^, showing therapeutic potential against cancers and neutropenia resulting from chemotherapy ^5–7^. Additionally, GM-CSF has been explored for cancer therapies as an immunocytokine, combined with antibodies or fragments, and fusokines, combined with other cytokines such as IL-2 and IL-4 ^8–10^.

GM-CSF is naturally glycosylated in mammalian cells with two *N*-linked glycosylation sites ^11^ and four *O*-linked glycosylation sites ^12,13^. The potential role of glycans in GM-CSF safety is highlighted by clinical studies that found that *Escherichia coli*-derived (*E. coli*-derived) GM-CSF elicits higher adverse reactions, such as the production of anti-GM-CSF antibodies, attributed to the absence of glycans ^14,15^. The degree and type of glycosylation also influence the properties of GM-CSF. Increased *N*-glycan site occupancy (two occupied *N-*glycosylated sites versus one) reduces GM-CSF receptor binding interactions ^11^ and decreases bioactivity, as demonstrated by reduced cell proliferation of bone marrow mononuclear cells ^16^ and AML193 cells, a human monocytic leukemia cell line dependent on GM-CSF ^11^. However, conflicting reports exist regarding the role of sialic acid. While one study suggests that sialic acid removal does not affect GM-CSF activity ^17^, another indicates that sialic acid enhances its *in vitro* bioactivity as measured by TF-1 cell proliferation, a GM-CSF-dependent human erythroleukemia cell line ^18^. In another study, researchers expressed GM-CSF glycovariants in human Namalwa cells, including forms with either only O-glycans (O) or both N- and O-glycans (2N+O). These glycoforms displayed distinct bioavailability profiles. GM-CSF variant with 2N+O glycans increases bioavailability by fivefold compared to both non-N-glycosylated (O) and *E. coli-*produced GM-CSF. This glycosylated form reached significantly higher concentrations in serum for a longer duration than non-glycosylated forms ^16^. These findings highlight the role of glycans in its activity and bioavailability.

Transcriptomic analysis of GM-CSF^+^ and GM-CSF^−^ human T cells isolated by cytokine secretion assay showed that MGAT5 was modestly yet highly significantly upregulated in GM-CSF-expressing versus non-expressing cells (log_2_ fold-change = 0.83; p = 9.77×10^-7^) ^19^. MGAT5 catalyzes β1,6-N-Acetylglucoseamine (GlcNAc) branching on N-glycans, which enables the formation of highly branched, galectin-binding N-glycans (N-acetyllactosamines) and thereby strengthens the galectin–glycoprotein lattice that regulates T-cell receptor clustering and activation thresholds for T cell activation ^20^. In another study, *Mgat5*-deficient mice displayed increased susceptibility to experimental autoimmune encephalomyelitis (EAE) and elevated levels of proinflammatory cytokines such as IFN-γ and TNF-α ^21^. Together, these findings make MGAT5 potentially a mechanistically and clinically relevant glycosyltransferase and provide motivation for studying how MGAT5 may modulate GM-CSF activity. Here, we investigated how the glycan structures of GM-CSF influence its activity. MGAT5 is expected to contribute to the glycan structure of GM-CSF, since it catalyzes the addition of a GlcNAc residue to the α1,6 mannose through a β1,6-linkage, which is important for forming tetra-antennary glycan structures ^22^. Because MGAT5-mediated branching may impact the degree of sialylation on proteins ^23^, we also investigated how changes in sialylation affected the protein. We hypothesized that elevated expression of branching glycosyltransferases, such as MGAT5, in GM-CSF–producing cells might play a functional role in enhancing protein activity. We produced multiple glycoforms using a panel of glycoengineered Chinese hamster ovary (geCHO) cells to obtain various glycoforms of GM-CSF and found that glycan microheterogeneity at the branching level impacts GM-CSF activity. Thus, future efforts to use GM-CSF or fusions thereof should carefully consider glycosylation as a modulator of activity.

## Results

### GM-CSF products vary in size in glycoengineered CHO cells

To study the impact of MGAT5 on GM-CSF glycosylation and activity, we expressed GM-CSF in CHO-S and a set of geCHO cell lines^24,25^. Specifically, six products, denoted GM-CSF-A to F (see Table 1 for the host cell genotypes), were purified and assessed via SDS-PAGE (Figure S1). We further subjected the products to anti-HPC4 western blot analysis. We found that GM-CSF from different samples had heterogeneous glycans spanning from 20 to 40 kDa. Wild-type (WT) and *Mgat5* knock-out (KO) were observed between 35-40 kDa. GM-CSF expressed in CHO cells engineered to remove α2-3-linked sialylation exhibited glycoforms spanning from 20 to 38 kDa, while GM-CSF with α2-3-linked sialylation predominantly appeared around 38 kDa, with a smeared band between ∼38-35 kDa. GM-CSF from cells also expressing human ST6GAL1 (adds α2-6-linked sialylation) and engineered to express bi-antennary glycans had more uniform glycoforms between 30-36 kDa, whereas the CHO cells engineered to enhance tetra-antennary structures showed more branched glycans, resulting in glycoforms between 30-40 kDa (Figure S2).

**Table 1.** Cell lines used in the study and their corresponding genotype.

| CHO/<br>geCHO ID | Expected sialylation | Expected most<br>abundant glycan<br>branching† | Knock-outs† | Knock-ins† | GM-CSF |
| --- | --- | --- | --- | --- | --- |
| CHO-S | α2,3-linked<br>sialylation | Mix of bi-, tri-<br>and/or<br>tetra-antennary <sup>26</sup> | - | - | **A** |
| CL107 | α2,3-linked<br>sialylation | Tri-antennary <sup>27</sup> | *Mgat5* | - | **B** |
| CL187 | No α2,3-linked<br>Sialic acids | Tetra-antennary | *St3gal3* + *St3gal4* +<br>*St3gal6* + *B3gnt2* + *Sppl3* | - | **C** |
| CL419 | α2,3-linked<br>sialylation | Tetra-antennary | *B3gnt2* + *Sppl3* | - | **D** |
| CL5957 | No α2,3-linked sialylation<br>+ α2,6-linked sialylation | Bi-antennary | *Mgat4a* + *Mgat4b* + *Mgat5* +<br>*St3gal3* + *St3gal4* + *St3gal6* +<br>*B3gnt2* + *Sppl3* | *hSt6gal1* | **E** |
| CL5967 | No α2,3-linked sialylation<br>+ α2,6-linked sialylation | Tetra-antennary | *St3gal3* + *St3gal4* + *St3gal6* | *hSt6gal1* | **F** |

In parallel, the corresponding producer cell lines were analyzed by RNA-Seq to evaluate whether GM-CSF overexpression altered global transcriptional programs. Differential expression analysis comparing GM-CSF–expressing geCHO cells with pUC19 vector controls identified only ∼1.5% of genes (FDR<0.05) as significantly differentially expressed, indicating that GM-CSF overexpression did not substantially perturb baseline cellular gene expression (Figure S3).

### geCHO-produced GM-CSF variants present diverse dominant glycan structures

Glycan analysis confirmed that variants produced in the geCHO cells, GM-CSF-D and GM-CSF-E, exhibited expected glycan branching patterns based on their host cell line genotypes (Table 2; host cell genotypes are listed in Table 1, and the expected most abundant glycan structures are listed in Table 3). GM-CSF-C predominantly displayed tri- and tetra-antennary glycans, with tri-antennary glycans being the most abundant. Interestingly, only 2.4% of GM-CSF-F glycans were identified as tetra-antennary, while 20.3% of the total peak area corresponded to unannotated peaks. Unannotated peaks may result from N-acetyllactosamine (LacNAc) extensions that can be generated due to *B3gnt2* expression ^27^ by GM-CSF-producer host cells (Table 1). Since such glycans with LacNAc extensions are not included in the glyXera database (glyXtoolCE™, glyXera GmbH, Germany), further analysis would be needed to confirm their presence.

**Table 2.** Glycoprofiling results of GM-CSF products with glycan branching information. The total glycoforms, categorized by their corresponding branching structures, are presented as the percentage of the total peak area (TPA). The glycoforms include mono-antennary, bi-antennary, tri-antennary, and tetra-antennary structures, with their respective distributions shown for each GM-CSF product.

| Host cell line expected sialylation profiles / GM-CSF product IDs | Mono-antennary structures (%TPA) | Bi-antennary structures (%TPA) | Tri-antennary structures (%TPA) | Tetra-antennary structures (%TPA) |
| --- | --- | --- | --- | --- |
| α2,3-linked sialylation (GM-CSF-A) | 14.4 | 17.5 | 19.8 | 32.4 |
| No α2,3-linked sialylation (GM-CSF-C) | 6.2 | 9.2 | 41.5 | 38.5 |
| α2,3-linked sialylation (GM-CSF-D) | 4.9 | 6.2 | 30 | 45.2 |
| No α2,3-linked sialylation and α2,6-linked sialylation-biantennary (GM-CSF-E) | 12.6 | 78.9 | 0 | 0 |
| No α2,3-linked sialylation and α2,6-linked sialylation-tetra-antennary (GM-CSF-F) | 29.7 | 26.5 | 21.2 | 2.4 |

**Table 3.** The expected most abundant glycan structures of the purified GM-CSF products.

| Product ID | Host cell line expected glycan sialylation profiles | Expected most abundant glycan structures |
| --- | --- | --- |
| GM-CSF-A | α2,3-linked sialylation (WT) | Mix of bi-, tri- and/or tetra-antennary, α2,3-linked sialylation |
| GM-CSF-B | α2,3-linked sialylation (MGAT5 KO) | Tri-antennary, α2,3-linked sialylation |
| GM-CSF-C | No α2,3-linked sialylation | Tetra-antennary; no α2,3-linked sialylation |
| GM-CSF-D | α2,3-linked sialylation | Tetra-antennary; α2,3 sialylation |
| GM-CSF-E | No α2,3-linked sialylation + α2,6-linked sialylation | Bi-antennary; no α2,3-linked sialylation; α2,6-linked sialylation |
| GM-CSF-F | No α2,3-linked sialylation + α2,6-linked sialylation | Tetra-antennary; no α2,3-linked sialylation; α2,6-linked sialylation |

In the glycan analysis, GM-CSF-E exhibited more homogeneous glycan branching, consisting only of mono- and bi-antennary structures (Table 2). It also showed more uniform band formation than other GM-CSF glycoforms in the SDS-PAGE analysis (Figure S1). In contrast, other samples, including GM-CSF-A, GM-CSF-C, GM-CSF-D, and GM-CSF-F, exhibited a broader range of glycan branching, including mono-, bi-, tri, and tetra-antennary structures. These glycovariants were detected as more pronounced smeared bands, which is common when multiple glycoforms of a protein are present (Table 2, Figure S1).

Glycan analysis with neuraminidase treatment also verified that the GM-CSF glycoforms exhibited the expected terminal sialic acid composition, as terminal sialic acids predominantly attached via α2,3 linkages in GM-CSF-A and -D and α2,6 linkages in GM-CSF-E and -F (produced from human *ST6GAL1* expressing cells)^28^, consistent with the host cell line glycosylation (Figure 1, Table 1). GM-CSF produced by WT CHO-S cells featured terminal sialic acids, with 41.7% linked via α2,3 linkage and 7.4% via α2,6 linkage. While inadequate amounts of GM-CSF-B were obtained to allow for glycoprofiling, its genotype indicates it should have sialylation patterns that mirror that of WT CHO-S cells, albeit with no tetra-antennary glycans. GM-CSF-C exhibited greatly reduced amounts of sialylation, with only 7.9% of glycans harboring terminal sialic acids linked via α2,3 linkage. In addition, GM-CSF-D exhibited 66.7% α2,3-linked terminal sialic acids. In contrast, GM-CSF produced by host cells expressing human *ST6GAL1* inserted host cells (CL5957 and CL5967, respectively) displayed only α2,6-linked terminal sialic acids (85.1% and 55.5%, respectively), as expected ^28^. These findings confirmed the sialic acid composition of our products and expected glycan branching, allowing us to proceed with testing their biological activity.

**Figure 1.**
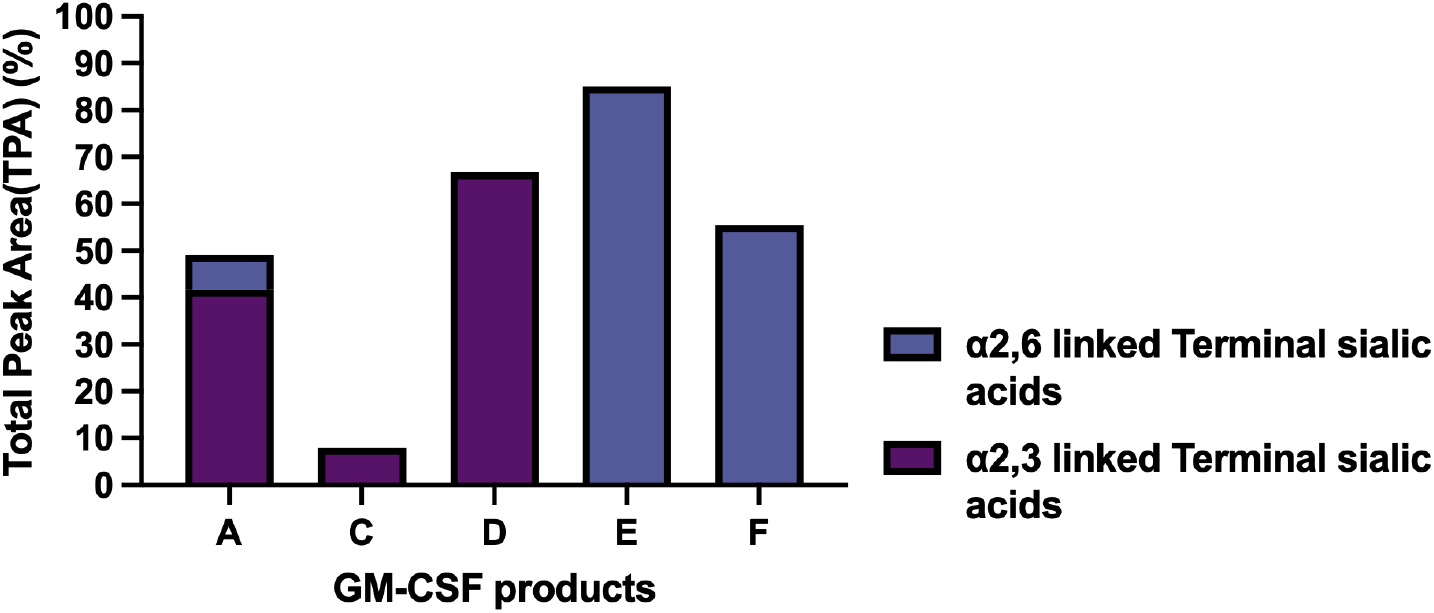
GM-CSF glycovariants exhibited expected terminal sialic acid abundances. Terminal sialic acid abundance regarding their linkage is shown as the percentage of total peak area (TPA%) for GM-CSF glycovariants: GM-CSF-A, -C, -D, -E, and -F.

### *Mgat5* removal decreases GM-CSF bioactivity

We assessed the biological impact of glycoengineered GM-CSF on TF-1 cell proliferation by comparing its activity to aglycosylated *E. coli*-produced GM-CSF (erh-GM-CSF) as a control. TF-1 cells, when treated with GM-CSF-A (produced in WT CHO cells), proliferated at a rate similar to those treated with erh-GM-CSF (Figure 2A and 2G).

**Figure 2.**
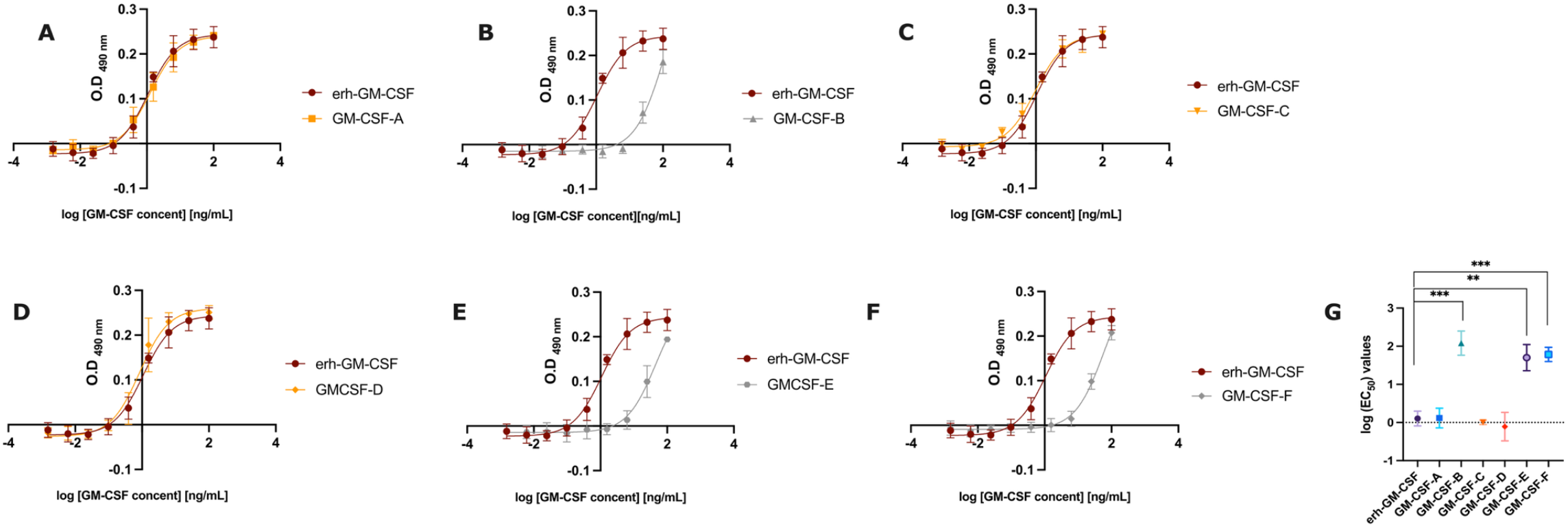
Dose-response curves showing the proliferation rate of TF-1 cells treated with GM-CSF glycovariants and a summary of their potencies. TF-1 cells were exposed to nine serial concentrations of each GM-CSF variant (0.003-100 ng/mL), and TF-1 proliferation was measured by optical density at 490 nm (OD_490_). The dose-response curves were generated using nonlinear regression analysis with a three-parameter logistic equation to obtain EC_50_ values (A-F). Mean log_10_-transformed EC_50_ values from three biological replicates are shown with error bars indicating the standard deviation (G). Statistical significance of GM-CSF bioactivity responses was assessed by comparing the log_10_-transformed values between the GM-CSF glycoforms and erh-GM-CSF using a two-tailed t-test. ** p < 0.01, *** p < 0.001.

Notably, GM-CSF derived from the *Mgat5* KO cell line (GM-CSF-B) and the human *ST6GAL1*-inserted cell lines (CL5957 and CL5967, corresponding to GM-CSF-E and GM-CSF-F, respectively) exhibited significantly reduced proliferation rates compared to erh-GM-CSF, with 20-, 16-, and 17-fold higher EC_50_ values, respectively, (Figure 2B, 2E, 2F, 2G; Table S1). GM-CSF-C and D (produced from CL187 and CL419 host cell lines) showed comparable activity to erh-GM-CSF, with similar EC_50_ values (Figures 2C-D, 2G; Table S1). However, GM-CSF-D exhibited variable EC_50_ values, potentially due to differences in biological responses among triplicates (Figure 2D, Table S2). Furthermore, GM-CSF-B, -E, and -F displayed significantly reduced bioactivity compared to GM-CSF derived from the WT cell line (GM-CSF-A) (Figure S4).

## Discussion

GM-CSF is a promising cytokine for treating immunosuppressed patients ^5^. This glycoprotein contains two *N-* and four *O-*glycosylation sites ^11–13^. Since glycosylation can be a critical quality attribute for therapeutic proteins, affecting their bioactivity and immunogenicity ^29^, considering the impact of glycosylation on the therapeutic use of GM-CSF is important. Prior research on GM-CSF has indicated that the removal of glycans can trigger immune responses by exposing GM-CSF epitopes. Therefore, glycans can reduce immunogenicity by masking these epitopes ^14^. However, the influence of branched glycans and sialic acids on bioactivity has not been clearly defined, and some conflicting results have been presented. Some studies suggest that the removal of sialic acids either does not affect ^17^ or reduces biological activity ^18^. In this study, we tested a panel of GM-CSF glycoforms to evaluate the impact of sialic acid presence and conformation, along with the influences of glycan branching on the protein’s biological activity.

Our study revealed that deleting the *Mgat5* gene in CHO cells, which can result in bi- and tri- antennary glycans ^27^, significantly reduced GM-CSF bioactivity. However, we were unable to assess this effect when α2,6-linked sialic acids (derived from CL5957 and CL5967 host cells with human *ST6GAL1* insertions) were present. Interestingly, GM-CSF-F displayed limited MGAT5 activity, with only 2.4% tetra-antennary glycans. GM-CSF-B and -E also lacked MGAT5 activity based on their host cell genotypes, and GM-CSF-F, exhibited low tetra-antennary glycan levels despite *Mgat5* expression. Since all three variants (B, E and F) showed high EC_50_ values, it seems the branch added by MGAT5 may significantly impact GM-CSF activity.

Moreover, we did not observe any impact of α2,3-linked sialic acids on GM-CSF bioactivity. Knocking out genes reported to be important for forming α2,3-linked sialic acids (CL187) and increasing the α2,3-linked sialic acid abundance (CL419) did not change the activity. This observation is consistent with earlier findings, which reported no differences in bioactivity, measured via radioimmunoassay and chronic myelogenous leukemia (CML) assay, between desialylated and WT CHO-derived GM-CSF ^17^.

However, our results contradict those of Hashimoto *et al*., who reported that desialylation reduced GM-CSF biological activity, as we did not observe increased GM-CSF bioactivity in sialylated CHO-derived GM-CSF glycoforms. Their study compared TF-1 cell proliferation levels among *E. coli*-, yeast-, and CHO-derived GM-CSF forms with or without sialidase treatment at a single concentration (15 pM) ^18^. Notably, they did not generate dose-response curves across varying concentrations, which would have allowed for a more comprehensive comparison of bioactivity levels. Therefore, we cannot interpret their results in the context of glycan structure.

We verified the sialic acid content of our products, which matched the host cell genotypes. Glycan analysis showed that WT GM-CSF harbored 7.4% α2,6-linked terminal sialic acids, consistent with previous studies showing that ST6GAL1, the enzyme responsible for adding α2,6-linked sialic acid to glycan structures, is reported to be silenced in CHO cells. ^30–32^. Small amounts of α2,6-linked sialic acid may have resulted from the low expression of *St6gal1* and 2. GM-CSF with no α2-3-linked sialylation exhibited 7.9% α2,3-linked terminal sialic acid; while the major ST3GAL isoforms needed for this linkage–*St3gal3, St3gal4*, and *St3gal6* ^27^–were deleted in the host cell line (CL187), three others—*St3gal1, St3gal2, and St3gal5*—are thought to add α2,3 linkages to O-glycans or glycolipids but have not been extensively studied in CHO cells ^33,34^. Thus, further research is needed to identify the source of this residual α2,3 sialylation.

Regarding *O-*glycans, GM-CSF has been reported to be *O*-glycosylated in CHO cells. However, the functional impact of these moieties has not been fully elucidated yet ^11–13,26^. In this study, we focused on the *N-*glycans due to the findings on *N*-glycosylation and observed gene co-expression of *N*-linked glycosyltransferases with GM-CSF across T cells ^19^, and we had access to a panel of *N*-glycoengineered CHO cell lines. Nevertheless, the potential influence of *O*-glycans on our results remains unexplored and warrants further investigation in future studies.

Glycoengineered cell lines with *B3gnt2* and *Sppl3* KOs enhanced the sialic acid abundance of our products. These genes are associated with competing sialic acid attachment and reduced glycosyltransferase prevalence in cells, respectively. B3GNT2 competes with sialyltransferases for the same substrate on tetra-antennary glycans to add poly-N-acetyllactosamine chains. The overexpression of *B3gnt2* has been linked to the suppression of sialylation, likely through reduced galactosylation, the sugar moiety to which sialic acids are typically attached ^35,36^. Furthermore, SPPL3 cleaves glycosyltransferases and decreases their abundance in the Golgi apparatus ^37^. These genes were deleted from the host cells producing GM-CSF-D (α2,3 sialylation glycoform) and GM-CSF-E (only α2,6 sialylation in biantennary form, derived from the host cell line with human *ST6GAL1* insertion, and deletion of *Sppl3* and *B3gnt2* and genes responsible for α2,3 sialylation, tri- and tetra-antennary structures). As expected, GM-CSF-D had more sialic acid than WT GM-CSF. GM-CSF-E also demonstrated higher sialic acid abundance compared to GM-CSF-F (only α2,6 sialylation glycoform, derived from the host cell line with a human *ST6GAL1* knock-in and KO of *St3gal3, St3gal4*, and *St3gal6*).

Glycans have been shown to reduce the production of antibodies against GM-CSF during clinical use ^14,15^ and determine the cytokine activity ^11,16^. Moonen *et al*. demonstrated that deglycosylation improved CHO-produced GM-CSF activity in CML and immunoassays ^17^. Human-derived GM-CSF variants with lower glycosylation levels (16-18 kDa and 23-25 kDa) displayed bioactivity comparable to erh-GM-CSF, whereas the highly glycosylated variant exhibited reduced activity ^11^. However, the role of sialylation in GM-CSF bioactivity has previously been studied only using sialidase-treated GM-CSF forms and comparing them with the non-sialylated forms, but earlier techniques were limited ^17,18^. Advances in cell line engineering techniques and detailed glycan analysis now allow for more comprehensive studies. Utilizing these tools, we are the first to investigate the role of glycan branching and both α2,3- and α2,6-linked sialic acids on GM-CSF bioactivity through glycoengineering.

We produced six GM-CSF glycoforms and glycoprofiled five, confirming their glycan structures, including sialic acid compositions. We highlighted the essential role of MGAT5 in maintaining GM-CSF bioactivity and observed that α2,3- and α2,6-sialylated variants exhibit differences in bioactivity (Figure 3). Because the α2,6-sialylated samples also differ in branching heterogeneity and may contain LacNAc extensions, the reduced activity cannot be attributed solely to the sialic acid linkage. Instead, the observed effects could arise from a combination of linkage, branching complexity, and LacNAc extensions. However, since we were unable to analyze the glycan structure of the product derived from the *Mgat5* KO host cell line, further glycan characterization is necessary to confirm structure-function relationships beyond branching associated with Mgat5.

**Figure 3.**
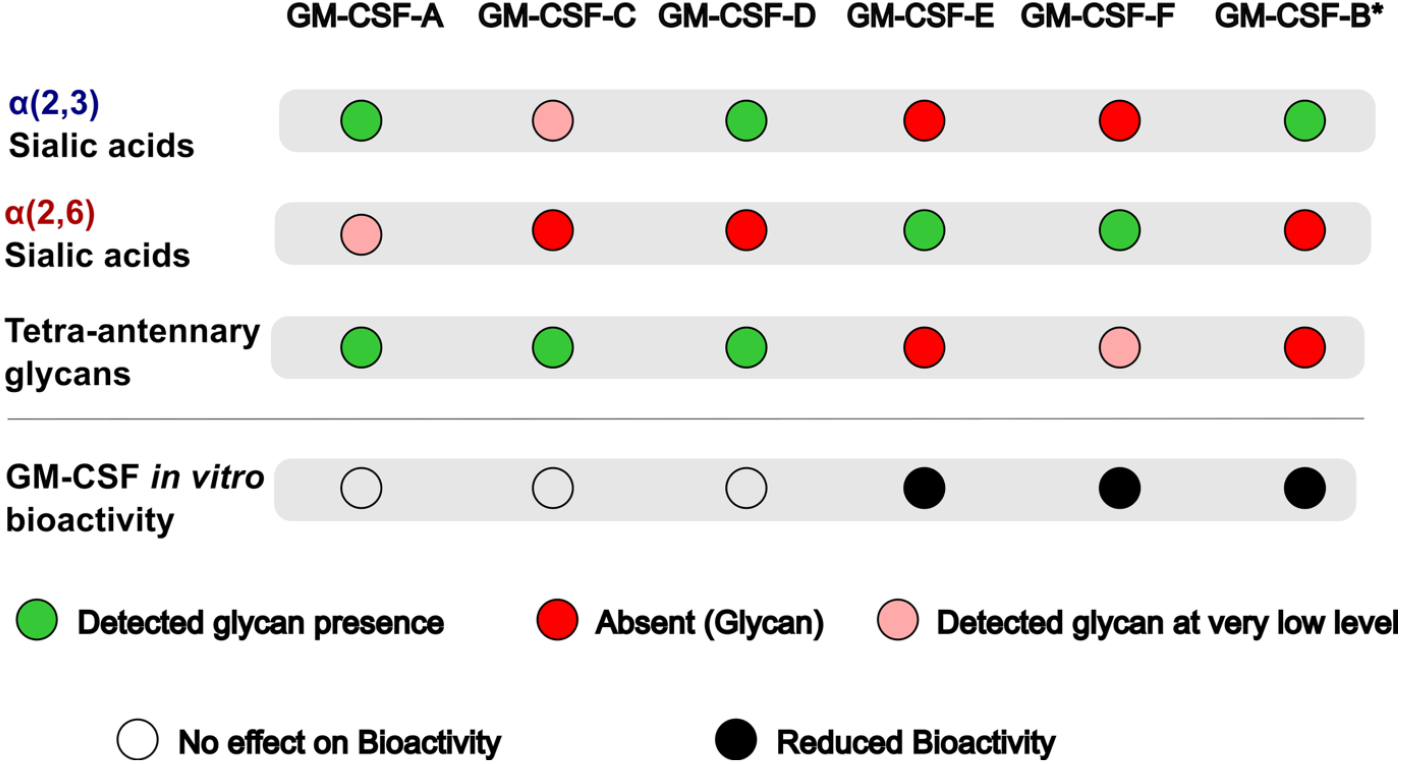
Summary of glycoforms of GM-CSF and their impact on *in vitro* bioactivity. The table illustrates sialic acid compositions (α2,3 or α2,6 linked), tetra-antennary glycan structures, and bioactivities of each GM-CSF glycoform. The asterisk (*) denotes the expected glycan structure of GM-CSF-B, based on an *MGAT5* KO cell line study ^27^. Green circles indicate the presence of a specific sialic acid form or tetra-antennary glycans, red circles indicate the absence of sialic acid moieties or tetra-antennary glycans. Light red circles show the detected glycans at very low levels. The grey circle denotes similar bioactivity levels, while the black circle indicates lower bioactivity compared to erh-GM-CSF.

In future studies, receptor-binding studies, as well as circular dichroism and mass spectrometry analyses, will be important to evaluate the impact of glycoforms and their conformational states on GM-CSF functionality. Additionally, the conformational change could be attributed to the reduced stability, as glycans can protect cytokines from protease degradation ^38,39^, heat denaturation ^40^, or oxidation ^41^. Therefore, these structural and stability effects should be taken into account when interpreting glycosylation-dependent changes in GM-CSF activity.

While research on GM-CSF bioactivity began in the late 1980s, much remains unknown about the impact of GM-CSF glycosylation. Nonetheless, GM-CSF has been engineered into fusokine and immunocytokine derivatives ^42^, and mammalian host cells have remained the preferred production systems. However, glycoengineered CHO cells produce altered glycan structures, which can significantly affect the functionality of the product. Therefore, careful characterization and control of glycan structures will be essential for the rational design of next-generation cytokine therapeutics.

## Material and Methods

### Plasmid design and cloning

An expression vector harboring GM-CSF with an HPC4 tag was generated with NEBuilder HiFi DNA assembly (NEB #M5520). The plasmid sequence was confirmed using Sanger sequencing (Eurofins) and extracted with Gigaprep kits (Macherey-Nagel, Cat#740548).

### Cell cultivation

CHO-S™ cells (Gibco, Cat#AT11364-01) were obtained, and glycoengineered cell lines were generously gifted by Bjørn Voldborg (DTU). Glycoengineered cells were generated as previously described (US20230272445A1), using the methods outlined by Rocamora *et al*. ^25^. Both wild-type and glycoengineered cell lines were used for GM-CSF production. The glycoengineered cell lines used in this study were generated through one or more KOs or knock-ins of glycosyltransferase genes, which are responsible for variations in *N*-linked glycan branching or sialylation (Table 1). The expected glycan structures resulting from the deletion or insertion of glycosyltransferases are illustrated in Figure 4 ^27^. However, the function of SPPL3 is not shown, as it does not add or remove specific glycan bonds. Instead, SPPL3 decreases glycosyltransferase abundance by cleaving them in the Golgi apparatus, which can, in turn, increase the glycan content of proteins ^37^.

**Figure 4.**
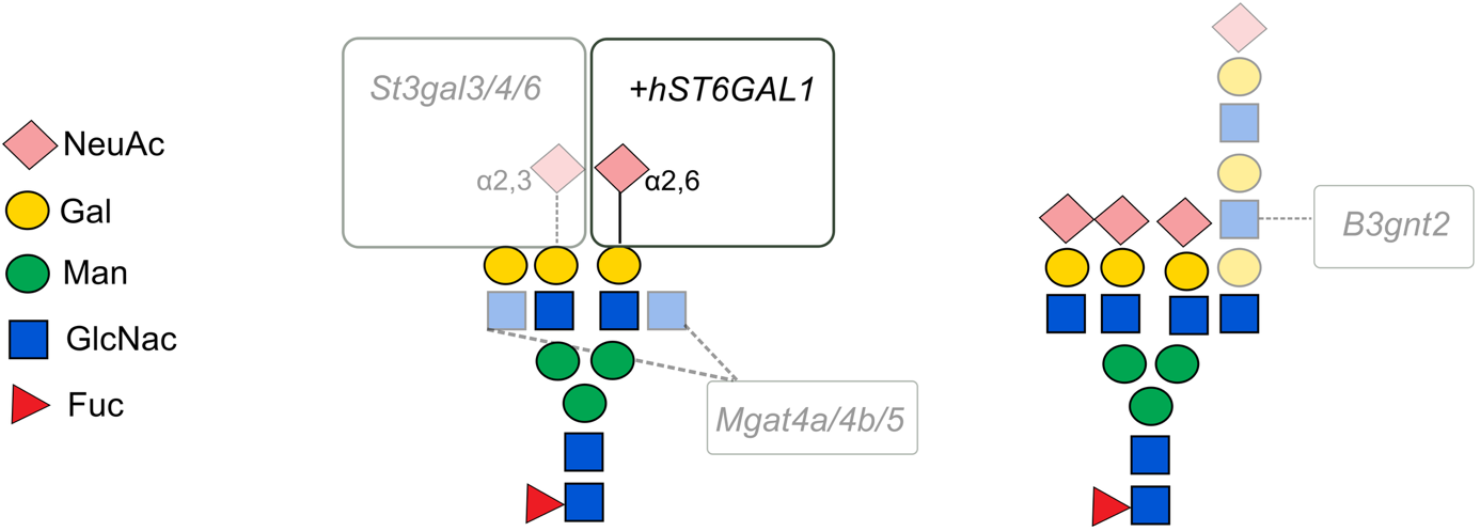
Illustration of tetra-antennary N-glycans capped with sialic acids (NeuAc) alone (left) and featuring a poly-N-acetyllactosamine extension of the β6 antenna and capped with sialic acids (NeuAc)(right). Lines indicate the reported functions of glycosyltransferase genes examined in this study ^27^. Transparent glycan moieties indicate their absence in cells with deletions of the genes listed in the grey box. The “+ *hST6GAL1*” represents the knock-in of that gene, resulting in the incorporation of a novel glycan moiety typically absent from CHO-produced proteins.

For cultivations, CD CHO medium (Gibco, Cat#10743029) supplemented with 8mM L-Glutamine (Gibco, Cat#A2916801) was used. Cells were maintained in 125 mL Polycarbonate Erlenmeyer shake flasks with vent caps (Corning, Cat #431143) at 37°C with 5% CO_2_ on an orbital shaker platform at 120 rpm with 25 mm amplitude. NucleoCounter NC-250 (ChemoMetec, Denmark) was used to monitor viable cells.

To test bioactivity, the TF-1 cell line (ICLC, Cat#HTL05001) was used, as it is a cell line derived from the bone marrow tissue of an erythroleukemia patient and is dependent on GM-CSF for cell maintenance. TF-1 cells were cultured in RPMI-1640 medium (Sigma-Aldrich, Cat#R5886-500ML) containing 2mM L-Glutamine (Gibco, Cat#A2916801), 20% FBS (Gibco, Cat#A5256801), and 5 ng/mL *E.coli-*derived GM-CSF (Peprotech, Cat#300-03). The cells were maintained in T25, T75, and T125 untreated flasks with vent caps (Corning, Cat#431463, #431464U, #431465). The flasks were incubated at 37°C with 5% CO_2_ without shaking. Cells were counted with AO/PI staining solution, prepared by a 1:1 mixture of acridine orange (Nexcelom, Cat#CS2-0106) and propidium iodide (Nexcelom, Cat#CS1-0109). For counting, 10 μL of the AO/PI staining solution and 10 μL of the cells were mixed and incubated for 30 seconds, and 10 μL of the solution was loaded on the cell counter (Denovix Celldrop FL). Viable cell densities were measured using an AO/PI assay.

### Transfection of GM-CSF and batch culture

One day before transfection, cells were seeded at 1.4 x 10^6^ cells/mL in 500 mL CD-CHO medium in 2L shake flasks (Corning, Cat#431255). The next day, transfection mixes were prepared with a final concentration of 1 μg/mL plasmid DNA in OptiPRO SFM media (Gibco) and 3.75 μg/mL LPEI MAX (Polysciences, Cat#24765-1). The cells were incubated at 37°C with 5% CO_2_ on an orbital shaker platform at 120 rpm as explained previously. On Day 1, cells were supplemented with 0.2% anti-clumping reagent and fed with 1% Tryptone N1 (Organotechnie, Cat#19553) and 0.6 mM Valproic acid sodium salt solution (Sigma, Cat#P4543-10g) to enhance recombinant protein production. Valproic acid inhibits the class I histone deacetylases (HDAC) and stimulates the proteasomal degradation of class II HDACs ^43^. Its supplementation after transient transfection has been shown to increase expression levels of antibody heavy chain and light chains by 10-fold and enhance antibody production by approximately fivefold in CHO DG44 cells ^44^. In addition, Tryptone N1, a peptone, has been reported to enhance recombinant protein production by boosting gene expression and translation in HEK293 cells when supplemented after 24 hours of transient transfection ^45^. Similarly, supplementing Tryptone N1 after transient transfection increased antibody yields in HEK293 cells ^46^. The cultures were incubated at 32°C, as described previously. On day 4, the supernatants were harvested through a two-step centrifugation process - first at 300 g for 5 minutes, and then the resulting supernatant was centrifuged at 1000 g for 10 minutes. The harvested materials were then stored at -70°C.

### Purification of GM-CSF products

Supernatants from a 500 mL cell culture batch, grown in 2L shake flasks (Corning, Cat#431255), were thawed overnight at 4°C and fully clarified by centrifugation at 4500 g for 30 minutes. Samples were concentrated fivefold by consecutive centrifugation using Amicon® Ultra Centrifugal Filters with a 3 kDa molecular weight cutoff (MWCO) (Sigma Aldrich, Cat#UFC900308) at 4500 g for 8 minutes at 4°C.

Purification was conducted on an ÄKTA pure system (GE Healthcare) with affinity chromatography using an anti-HPC4 (anti-protein C) tag antibody matrix (Roche, Cat#11815024001). The matrix was packed in a 2mL HiScale(Cytiva) column, and the purification method relies on the interaction between the HPC4 antibody and the Protein C-tag. This reaction is calcium-dependent, requiring the addition of 1mM CaCl_2_ to the harvests. Buffers were prepared according to the manufacturer’s instructions. The running method is explained in Table S3. The column was regenerated to discard all non-bound proteins, and the proteins were eluted as 1 mL fractions into a 96-deep well plate (Corning, Cat#CLS3960).

After purification, products were concentrated for bioactivity and glycan analysis by sequential centrifugation using Amicon® Ultra Centrifugal Filters with the same cutoff (Sigma Aldrich, Cat#UFC8003) at 4500 g for 8 minutes at 4°C. The goal was to achieve a concentration range of 0.2-0.5 mg/mL for glycan analysis.

### Verification of GM-CSF production with Western blot

First cell harvests were prepared for SDS-PAGE electrophoresis. Cell harvests were loaded as 13 μL sample along with NuPAGE^TM^ LDS Sample Buffer (Invitrogen, Cat#NP0322BOX) and loaded onto NuPAGE^TM^ 4-12% Bis-Tris Protein Gels (Invitrogen, Cat#NP0322BOX). The SDS-PAGE gels were run at 200V, 120 mA for 35 minutes.

Next, proteins were transblotted to iBlot nitrocellulose miniTransfer Stacks (Invitrogen, cat#IB301002) following the manufacturer’s instructions using the iBlot 2 Gel Transfer Device (Invitrogen, Cat#IB21001) with the P3 (20V, 5 minutes) program. Membranes were blocked with 5% (w/v) skim milk powder in PBS-T solution for 1 hour, then incubated with primary antibodies corresponding to the overexpressed proteins overnight at 4°C on the rotator. Membranes were washed three times for 5 minutes each with PBS-T. Next, samples were incubated with secondary antibodies for 1 hour at room temperature. GM-CSF proteins were detected by using Protein C-Tag antibody (HPC4) (GenScript, Cat#A00637-40, 1:1000) as the primary antibody and Goat Anti-Rabbit IgG antibody (Abcam, Cat#ab6721, 1:3000) as a secondary antibody. Membranes were rewashed as described and then incubated with 1 mL AmershamTM ECL^TM^ Prime Western Blotting Detection Reagent (GE Healthcare, Cat#RPN2232). Membranes were visualized using a chemiluminescent setting and 1 second exposure time on Amersham Imager 600 (GE Healthcare).

### Determination of the purity of purified products with SDS-PAGE

Purified products (2 μg per sample) were prepared with NuPAGE™ LDS Sample Buffer and analyzed by SDS-PAGE under the same conditions as described above. Gels were stained with Coomassie Brilliant Blue R-250 and imaged using the Amersham Imager 600 (GE Healthcare).

### TF1 assay

TF-1 cells were subjected to starvation by passaging them in media that did not contain GM-CSF and then incubated for 24 hours. The next day, cells were washed by centrifuging at 300 g for 10 minutes and then counted. The cells were resuspended in fresh media without GM-CSF, and 50,000 cells (0.1 mL per well) were seeded into each well of a 96-well plate (Corning Costar, Cat#3595). Recombinant erh-GM-CSF (Gibco PeproTech®, Cat#300-03-100UG) and the glycovariant products were diluted to concentrations of 100, 25, 6.25, 1.56, 0.39, 0.10, 0.024, and 0.002 ng/mL through serial dilution in GM-CSF-free culture media. The plate setup was designed with triplicates for each condition, with a total volume of 200 μL per well. GM-CSF-free media was added to blank wells, and erh-GM-CSF and glycoengineered product dilutions were added to the wells for GM-CSF treatment. Plates were incubated for 72 hours, and viable cell proliferation was monitored using CellTiter 96® AQueous One Solution Cell Proliferation Assay (Promega, Cat#3582), which contains MTS (3-(4,5-dimethylthiazol-2-yl)-5-(3-carboxymethylphenyl)-2-(4-sulfophenyl)-2H-tetrazolium) tetrazolium inner salt. The MTS solution is reduced by mitochondrial enzymes, generating formazan salt with a color change. After 3 hours of incubation, the plate was read at 490 nm (MTS) signal and 630 nm background by a plate reader (BMG LabTech).

### Glycan analysis

Glycan analysis was outsourced to Glyxera. First, *N*-glycan moieties were released with the PNGase enzyme, and *N*-glycome was further treated with two neuraminidases with different specificities: Sialidase A(α2-3, 6, 8, 9) and Sialidase S(α2-3). The glycan structures were analyzed by multiplexed capillary gel electrophoresis (CGE) with laser-induced fluorescence detection (xCGE-LIF) based Standard GlycoprofilingPLUS ^47^. This method enables the characterization of mono-, bi-, tri-, and tetra-antennary *N*-glycans and the composition and linkage specificity of sialic acids.

### Statistical analysis

Nonlinear regression analysis was employed to generate a dose-response curve model with a three-parameter logistic equation using GraphPad Prism (version 10). EC_50_ values, which represent the agonist concentration that gives a response halfway between the maximum and minimum responses, were calculated for each GM-CSF glycovariant using the following equation, where *X* represents the logarithms of GM-CSF concentrations, *Y* represents the absorbance value, and *Maximum* and *Minimum* represent the top and bottom absorbance values:

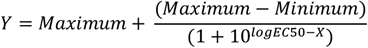

The EC_50_ was calculated from each biological triplicate, log_10_-transformed, and compared between groups using a two-sided t-test. A p-value of <0.05 was considered statistically significant.

The workflow of the study, from vector generation to producing and analyzing glycovariants, is depicted in Figure 5.

**Figure 5.**
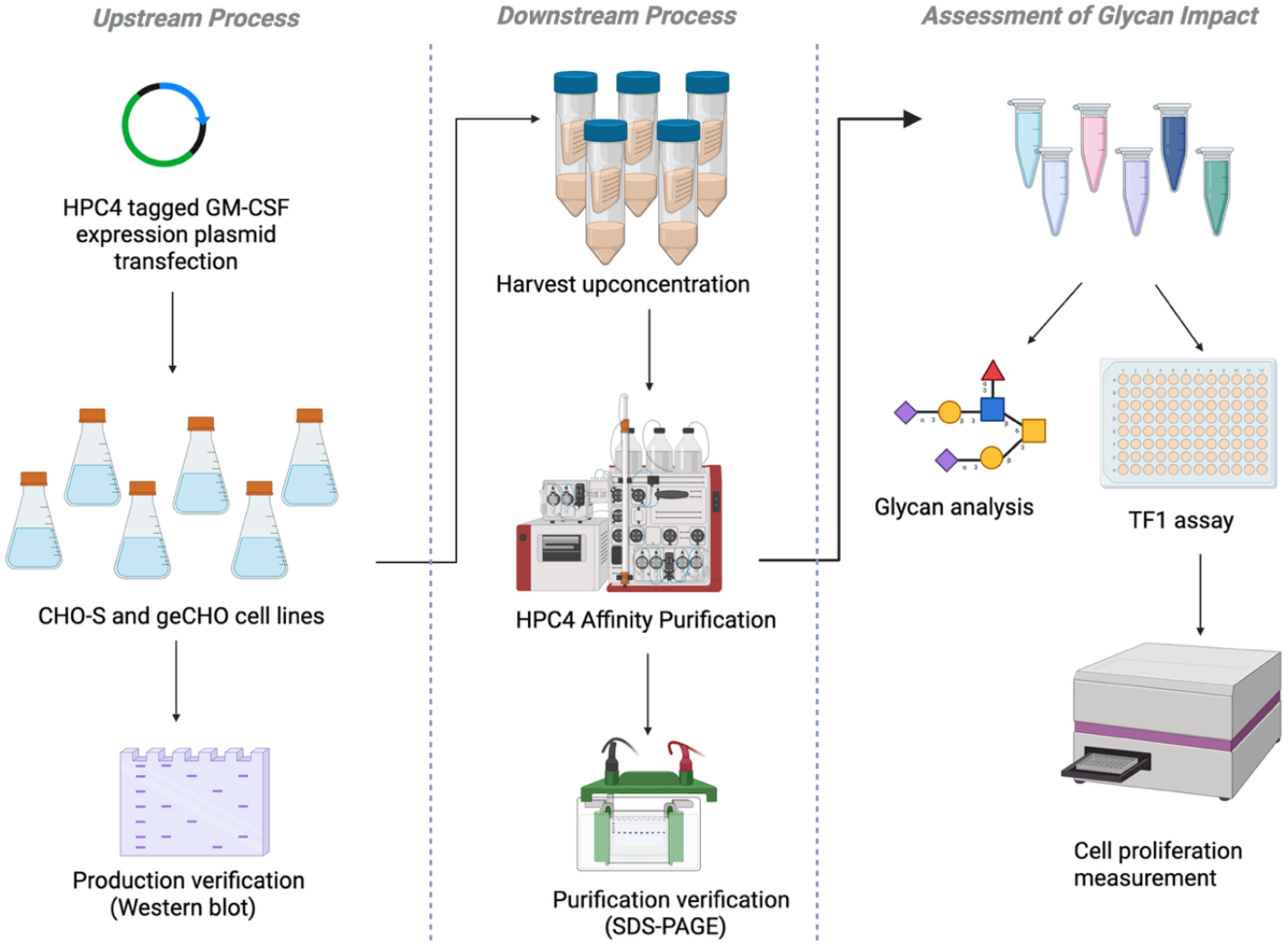
Study flowchart. First, an HPC4-tagged GM-CSF expression vector was generated and transiently transfected into glycoengineered CHO cell lines to produce various GM-CSF glycovariants. The successful production of multiple glycoforms was confirmed by western blot. The glycovariants were then purified using HPC4 affinity chromatography. After verifying product purity, glycan profiling was performed, and GM-CSF bioactivity was assessed using a TF-1 assay. Created with BioRender.com

## Acknowledgements

This work was supported by generous funding from National Institutes of Health (R35 GM119850) to NEL and Novo Nordisk Foundation (NNF20SA0066621) to NEL, SG and LKN. SG additionally received support from NNF19SA0056783, NNF21SA0072683, and NNF19SA0035474. LKN was supported by CFB2.0 Grant (NNF20CC0035580) and NNF14OC0009473, and NP was supported by NNF21OC0071624.

## Author contributions

EC designed and carried out experiments and data analysis. EC and NEL wrote the manuscript. SLS and JPA contributed to bioactivity assay development, and JPA contributed to bioactivity assay experiments. BGV, LKN, and SG provided critical feedback on research methods. LAD and HH aided in RNA-Seq analysis. SS and NP contributed to the purification design, and GS and KSF contributed to the purification experiments. All authors reviewed and provided critical feedback on the manuscript.

## Generative AI statement

During the preparation of this work ChatGPT and Perplexity were used to improve clarity and language flow. After using this tool, the authors reviewed and edited the content as needed and took full responsibility for the publication content.

