## Supplementary Information for "The Role of Glycan Structures in Modulating GM-CSF Bioactivity: Insights from Glycoengineering"

**Supplementary Figures**

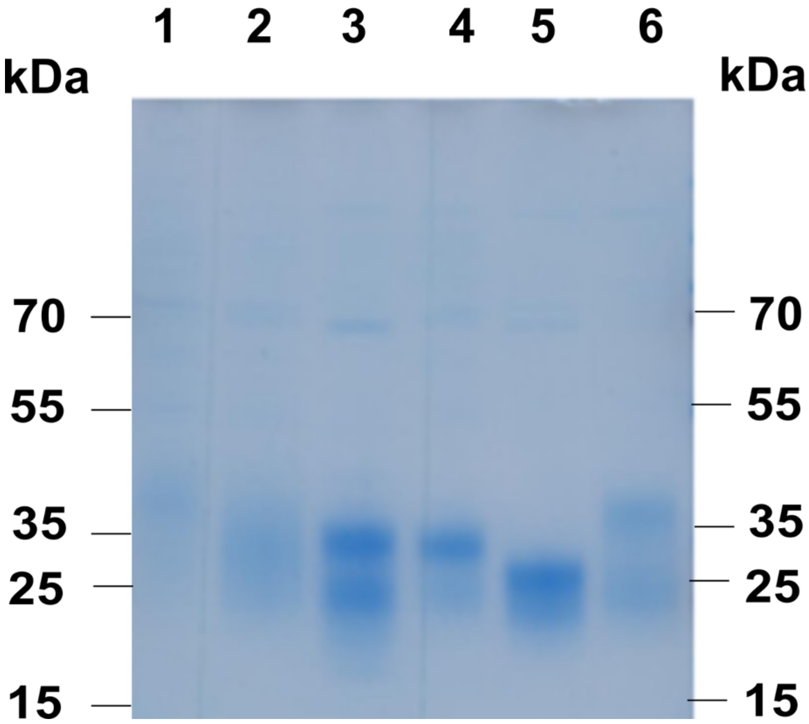


**Figure S1:** SDS-PAGE image for purified GM-CSF products (1: wild-type (GM-CSF-A), 2: *MGAT5* KO (GM-CSF-B), 3:No α2-3-linked sialylation (GM-CSF-C), 4: α2-3-linked sialylation (GM-CSF-D), 5:α2-6-linked sialic acids-bi-antennary form (GM-CSF-E), and 6: α2-6-linked sialic acids-tetra-antennary form (GM-CSF-F)). The observed bands corresponded to those detected in the western blot of the harvest samples (Figure S2).

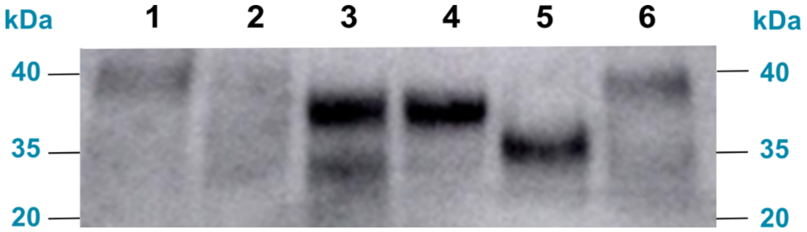


**Figure S2:** Western blot image of harvests from CHO-S and glycoengineered cell lines (1-GM-CSF-WT: CHO-S, 2-GM-CSF-*MGAT5* KO: CL107, 3-GM-CSF-no α2-3-linked sialylation: CL187, 4-GM-CSF-α2-3-linked sialylation: CL419, 5-GM-CSF-α2-6-linked sialic acids-biantennary structure: CL5957, 6-GM-CSF-α2-6-linked sialic acids-tetraantennary structure: CL5967).

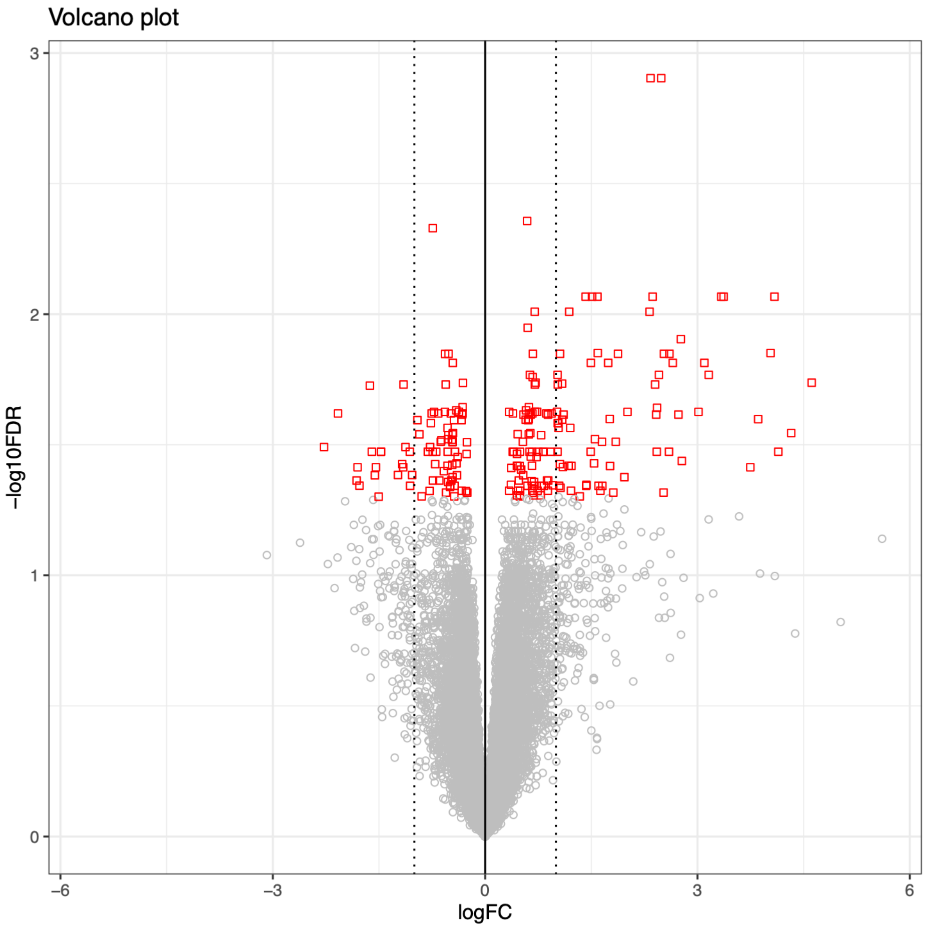


**Figure S3:** Volcano plot showing differential gene expression in geCHO cell lines comparing GM-CSF expression vs pUC19 controls. Each point represents a gene; $\log_2$ fold-change (logFC) displayed on the x-axis and statistical significance expressed as ($-\log_{10}$FDR) on the y-axis. Genes are color-coded by FDR: red (FDR $\leq 0.10$) and and grey (FDR $> 0.10$). Vertical dotted lines indicate $\pm 1$ $\log_2$ fold-change. No explicit fold-change cutoff was applied for significance.

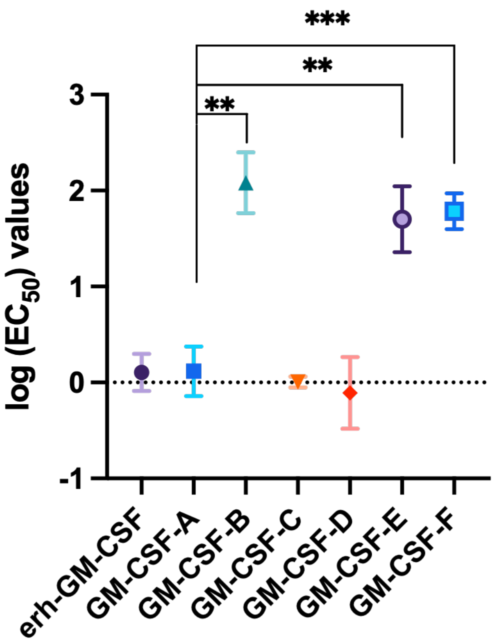


**Figure S4:** Log-transformed EC50 values (base-10) calculated from nonlinear regression analysis using a three-parameter equation across GM-CSF products. The data represent the mean log-transformed EC50 values from three biological replicates, with error bars indicating standard deviation. The statistical significance of GM-CSF bioactivity responses was assessed by comparing WT GM-CSF using a t-test. ** $p < 0.01$, *** $p < 0.001$.

### Supplementary Tables

**Table S1:** Mean of $\log_{10}$-transformed $EC_{50}$ values for each GM-CSF glycoform across three biological triplicates (Table S2) calculated by nonlinear regression analysis using a three-parameter equation.

| GM-CSF glycoforms | erh-GM-CSF | A | B | C | D | E | F |
|---|---|---|---|---|---|---|---|
| EC50 values | 0.11 | 0.12 | 2.08 | 0.01 | -0.11 | 1.70 | 1.79 |

**Table S2:** Log-transformed EC50 values (base-10) calculated from nonlinear regression analysis using a three-parameter equation for GM-CSF variants. Data represent individual values from triplicate experiments.

| GM-CSF variants | EC50-1 | EC50-2 | EC50-3 |
|---|---|---|---|
| erh-GM-CSF | 0.048 | 0.320 | -0.053 |
| GM-CSF-A | 0.197 | 0.327 | -0.174 |
| GM-CSF-B | 1.859 | 2.443 | 1.941 |
| GM-CSF-C | -0.053 | 0.058 | 0.010 |

**Table S2:** (continued)

| | | | |
|---|---|---|---|
| GM-CSF-D | -0.141 | 0.282 | -0.466 |
| GM-CSF-E | 2.042 | 1.705 | 1.355 |
| GM-CSF-F | 1.990 | 1.622 | 1.746 |

**Table S3:** HPC4 affinity chromatography running method

| Step name | Sample/buffer | Flow rate | Duration |
|---|---|---|---|
| Sample application | Sample | 0.5 mL/min | Depends on sample volume |
| Column wash | Equilibration/ Wash buffer | 1 mL/min | 15 CV |
| Elution | Elution buffer | 0.25 mL/min | 5 CV |
| Regeneration | Regeneration buffer | 1 mL/min | 10 CV |
| Equilibration | Equilibration/ Wash buffer | 1 mL/min | 10 CV |

### Supplementary Methods

#### RNA-seq analysis

***Cell cultivation and transfection:*** For the batch culture study used for RNA-Seq analysis, CHO-S and geCHO cell lines were cultured as explained in the "Cell cultivation" subsection of the Methods. These cell lines were seeded at 1.4 x $10^6$ cells/mL in 30 mL CD-CHO medium in 125 mL shake flasks (Corning, Cat #431143) and were transfected with plasmids expressing HPC4-tagged GM-CSF, AAT, and pUC19 separately. On Day 1, the same supplementation was performed with 0.2% anti-clumping reagent, 1% Tryptone N, and 0.6 mM Valproic acid sodium salt solution, and the cultures were subsequently incubated at 32°C. On days 3, 4, and 5, approximately 3 x $10^6$ cells were harvested, centrifuged at 300 g for 10 minutes, and the cell pellet was resuspended in 350 µL RLT/DTT buffer (Qiagen, Cat#79216) and stored at -80°C.

***Sample preparation:*** RNA was extracted from the cell lysates using the RNeasy Mini Plus Kit (Qiagen, Cat#74134), following the manufacturer's instructions. RNA concentration was measured using a Qubit fluorometer (Thermo Fischer Scientific), and the quality was assessed with a Fragment Analyzer system (Agilent). Preparation of RNA library and transcriptome sequencing were performed by Novogene Co., Ltd (UK) using the Illumina NovaSeq 6000 Sequencing platform.

***Data processing and quality control:*** The *Cricetulus griseus* reference transcriptome (rna.fna.gz) for release 104 was retrieved from NCBI in fasta format (RefSeq accession 20140878; GenInfo accession 7269151; RefSeq assembly GCF_003668045.3; GenBank assembly GCA_003668045.2; Name CriGriPICRH-1.0). Sample metadata were parsed from a design table describing experimental conditions (*GM-CSF* expression vs no *GM-CSF* expression) and used to construct the design matrix for differential expression analysis. Raw paired-end FASTQ reads were assessed for quality using FastQC (v0.12.1) [1]. Adapter and low-quality bases were removed using Trimmomatic (v0.39) [2] in paired-end mode. Transcript quantification was performed using kallisto (v0.50.1)[3] in pseudo-alignment mode with 100 bootstrap replicates and 8 threads. Following alignment, expression matrices estimating transcripts per million (TPM), counts per million (CPM), and averaged transcript length across samples were retrieved via tximport [4], applying the countsFromAbundance 'lengthScaledTPM' and 'no' as commands, respectively.

For data transformation and visualization, count matrices were then analyzed using EdgeR [5], DESeq2 [6], and limma [7]. Library sizes were calculated prior to filtering to confirm comparable sequencing depth across samples. Lowly expressed genes were removed using the filterByExpr function from edgeR [8], requiring a minimum of ten counts in at least 70 % of samples. Library sizes before and after filtering were compared to verify minimal data loss due to the removal of low-count records.

Normalization was performed using multiple methods to evaluate consistency across transformations: log-transformed raw counts (log(count+2)), transcript length–scaled TPMs, trimmed mean of M-values (TMM)-adjusted counts (*edgeR*), relative log expression (RLE) normalization (*DESeq2*), and $\log_2$-counts per million (logCPM) from *limma-voom*. Normalized expression distributions were visualized using density, box, and violin plots to assess inter-sample variability and confirm the effectiveness of normalization. PCA of all normalized expression matrices confirmed consistent clustering patterns across methods. Alignment, QC, and read count quantification were performed in R (v4.3.3)

***Differential gene expression analysis:*** Differential gene expression analysis was performed to determine the transcriptional impact of GM-CSF overexpression in CHO cell lines. "Experimental" samples comprised five independent geCHO cell lines (CL107, CL187, CL419, CL5957, CL5967) each transiently expressing GM-CSF, as well as wild-type CHO-S cells subjected to GM-CSF transfection (see Table 1 for genotypes). "Control" samples consisted of wild-type CHO-S parental lines overexpressing the pUC19 empty vector. All samples were collected in biological replicates across three time points (days 3, 4, and 5), yielding a total of six "Experimental" and three "Control" replicates.

Differential expression was assessed using the edgeR package with glmQLFit and glmQLTest functions [9,10], which model count data using a negative binomial distribution and estimate gene-specific dispersion to account for biological variability. The quasi-likelihood F test (glmQLTest) was used to compare global transcriptomic profiles between "Experimental" and "Control" groups.

For model specification, the categorical variable "Condition" was defined with two levels: "Experimental" (GM-CSF overexpression) and "Control" (vector-only CHO-S). When using DESeq2, "Control" was set as the reference level and "Experimental" as the contrast, such that effect sizes ($\log_2$ fold-changes) quantify expression differences in GM-CSF-overexpressing samples relative to vector-only controls. Differentially expressed genes were defined by an adjusted p-value of $< 0.05$ (Benjamini-Hochberg procedure), no log fold change cut-off was applied.
